# Endomucin regulates the endothelial cytoskeleton independent of VEGF

**DOI:** 10.1101/2024.07.17.603969

**Authors:** Jean Moon, Suman Chaudhary, Lorena Rodriguez Martinez, Zhengping Hu, Patricia A. D’Amore

**Affiliations:** Schepens Eye Research Institute of Massachusetts Eye and Ear, Boston, MA, USA; Department of Ophthalmology, Harvard Medical School, Boston, MA, USA; Department of Pathology, Harvard Medical School, Boston, MA, USA

## Abstract

The endothelial glycocalyx, lining the apical surface of the endothelium, is involved in a host of vascular processes. The layer contains a network of membrane-bound proteoglycans and glycoproteins. One such glycoprotein is endomucin (EMCN), which our lab has revealed is a modulator of VEGFR2 function. Intravitreal injection of siEMCN into the eyes of P5 mice impairs vascular development. In vitro silencing of EMCN suppresses VEGF-induced proliferation and migration. Signaling pathways that drive cell migration converge on cytoskeletal remodeling. By coupling co-immunoprecipitation with liquid chromatography/mass spectrometry, we identified interactions between EMCN, and proteins associated with actin cytoskeleton organization. The aim of the study was to investigate the influence of EMCN on cytoskeleton dynamics in angiogenesis. EMCN depletion resulted in reduction of F-actin levels, whereas overexpression of EMCN induced membrane protrusions in cells that were rich in stress fibers. The reorganization of the actin filaments did not depend on VEGFR2 signaling, suggesting that EMCN is a potential connection between the cytoskeleton and the glycocalyx.

## 1. Introduction

The luminal surface of endothelial cells (EC) is lined by a sugar-rich layer known as the glycocalyx, which forms the interface between the bloodstream and vascular endothelium [1, 2]. The glycocalyx participates in the regulation of vascular permeability, leukocyte adhesion and migration, mechanosensation and transduction, and bioavailability of angiogenic factors [3–6]. The basic structure consists of glycoproteins bearing sialic acids and proteoglycans decorated with glycosaminoglycan (GAG) chains. Heparan sulfate, chondroitin sulfate, and hyaluronic acid are the predominant GAGs that impart an overall negative charge, reinforcing the glycocalyx as an electrostatic barrier [6]. In addition to the membrane-bound proteins, growth factors noncovalently bind to the GAGs and adsorbed proteins such as albumin collectively contribute to vascular homeostasis [4, 7, 8].

The glycocalyx serves as a protective layer by repelling molecules and preventing leukocyte adhesion [9, 10]. However, it also facilitates interactions with extracellular ligands such as growth factors, thereby enabling proper signaling. Among the most characterized ligands is vascular endothelial growth factor (VEGF), which plays a critical role in physiological and pathological angiogenesis [11, 12]. Pro-angiogenic signaling, mediated by the VEGF/VEGF receptor 2 (VEGFR2) system, involves EC proliferation and migration [12–14]. Several studies identified endomucin (EMCN), a membrane-bound glycocalyx glycoprotein, as a key regulator of VEGFR2 internalization and downstream signaling. The knockdown of EMCN suppresses VEGF-induced proliferation, migration, and tube formation in human retinal endothelial cells (hREC) [15–18].

During angiogenesis, rapid proliferation and motility of ECs is achieved by alterations in actin cytoskeleton dynamics. The remodeling of actin underlies the shift from quiescence to angiogenic endothelial cells [19, 20]. In the quiescent state, the actin forms a cortical actin rim that undergoes consistent remodeling through polymerization and depolymerization [21]. Cells form protrusions for migration, and the glycocalyx has been reported to influence bending membrane bending [22–24]. Structural evidence indicates that the glycocalyx is anchored to the filamentous actin [25].

Though prior findings indicate that the glycocalyx, plasma membrane, and actin network are coupled and behave as a functional unit, the nature of that interaction remains unidentified [26, 27]. Our results suggest that EMCN forms complexes with actin binding proteins and regulators. We have shown that silencing of EMCN affects the formation of F-actin stress fibers. In this study, we examined the role EMCN has on F-actin levels and the cell morphology. Furthermore, we characterized the contribution that VEGFR2 has on the relationship between EMCN and absolute levels of F-actin.

## 2. Methods and Materials

### 2.1 Ethics Statement

All animal procedures were approved by the Harvard Medical Area Standing Committee on Animals and Institutional Animal Care and Use Committee of the Schepens Eye Research Institute/Mass Eye and Ear. Mice were handled in accordance with the National Institute of Health Guide for the Care and Use of Laboratory Animals.

### 2.2 Reagents and antibodies

Control siRNA (siCtrl, D-001810-01-05), siRNA directed against EMCN (siEMCN, L-015860-01-0005), and siRNA directed against VEGFR2 (siVEGFR2, L-003148-00-0005) were purchased as Smartpools (Dharmacon, Lafayette, CO, USA). Dharmafect 1 transfection reagent (T-2001-02; Dharmacon) was used for the in vitro studies. paraformaldehyde (32% PFA, 15714-S) was purchased from Electron Microscopy Sciences and diluted to 4% PFA in phosphate buffered saline (PBS). Phalloidin conjugated to Alexa Fluor 594 (A12381) was purchased from Thermo Fisher Scientific. dynamin I/II (2342; 1:1000), β-tubulin (2146; 1:1000), mouse anti-Myc (9B11; 1:1000) purchased from Cell Signaling were diluted in 3% bovine serum albumin (BSA)-PBS. Rat EMCN antibody (sc-65495) was purchased from Santa Cruz. 4X Laemmli’s SDS Sample Buffer (BP-110R) was purchased from Boston BioProducts (Ashland, MA, USA). Aflibercept (Regeneron Pharmaceuticals) was used at 1 nM. Adenovirus of myc-tagged full length human EMCN (Ad-hEMCN), GFP (Ad-GFP), and Cre recombinase were ordered from VectorBuilder (Chicago, IL, USA).

### 2.3 Mice

Floxed EMCN mice were designed and generated with Cyagen Bioscences. LoxP sites were inserted into the EMCN allele flanking intron 1 and 10. A GFP cDNA cassette was knocked in, and expression is under control of the EMCN promoter upon Cre recombination. Founder EMCN^flox/wt^ mice were bred to generate homozygous floxed mice.

### 2.4 Choroidal explant sprouting assay

Adult mice (6-8 wk-old) were euthanized, and their eyes were enucleated and kept in cold PBS on ice before dissection. The cornea and the lens were removed from the anterior of the eye, and the choroid and sclera from the retina. The choroid was cut into 1 mm^2^ pieces. Choroidal fragments were placed in 30 ul cold growth factor-reduced Matrigel™ (Trevigen) seeded in 24-well plates. After seeding the tissues, the plates were incubated in a 37°C cell culture incubator for 30 min to solidify the gel. EGM-2 BulletKit medium (500 ul) (Lonza, #CC-3162) supplemented with 5% fetal bovine serum (FBS) (Atlanta Biologicals), 2 mM L-glutamine (Lonza, #17-605E), and 100 U/mL penicillin–100 μg/mL streptomycin (Lonza, #17-602E) was added to each well and incubated at 37°C with 5% CO2 for 48 hr. Medium was changed every 48 hr.

Adenovirus-Cre was introduced between day 3 and 4. Phase contrast photos of individual explants were taken daily using EVOS2 microscope between days 3 to 6. The areas of sprouting were quantified using ImageJ. Explants were fixed in 4% PFA, permeabilized using 0.5% Triton-X 100, and stained for EMCN. Immunofluorescence images were obtained using EVOS2 microscope.

### 2.5 Cell Culture

Primary hRECs (P3) were purchased from Cell Systems (Troisdorf, Germany) and were cultured on glass cover slips that were coated with 0.2% gelatin-PBS for 30 min at 37°C. hRECs were cultured in EGM-2 BulletKit medium (Lonza, Basel, Switzerland, #CC-3162) supplemented with 2% fetal bovine serum, 2 mM L-glutamine (Lonza, #17-605E), and 100 U/mL penicillin–100 μg/mL streptomycin (Lonza, #17-602E) and were maintained in incubator at 37 °C with 5% CO_2_.

### 2.6 Immunoprecipitation and mass spectrometry

hRECs were plated on 150 mm dishes and cultured in complete EBM-2 media. Cells were collected in cold PBS and centrifuged at 1,500 rcf for 5 min at 4 °C. The pellet was resuspended in lysis buffer supplemented with protease inhibitors for 30 min on ice. The lysate was centrifuged at 16,000 rcf at 4 °C, and the supernatant was incubated with a mouse anti-Myc antibody (1:25), rotating overnight at 4 °C. Protein A/G Beads (Santa Cruz) were washed with 0.2% PBS and beads were blocked in 3% BSA, 0.5% PBST at 4 °C overnight. The beads were washed in PBS and incubated with lysate overnight, and the proteins were eluted using Laemmli’s SDS sample buffer with 100 mM DTT and incubation at 95 °C for 10 min. The proteins were separated on a 5% SDS-PAGE gel for 30 min at 60 V. The gel was stained with Coomassie blue and the lane excised. Gel sections were sent to Harvard Medical School Taplin Mass Spectrometry Core Facility for liquid chromatography with tandem mass spectrometry analysis. The annotated EMCN-specific binding proteins were further analyzed using PANTHER protein database for further functional clustering.

### 2.7 siRNA knockdown

siEMCN (50 nM, L-015860-01-000), control siRNA (siControl) (50 nM, D-001810-01-05), or siVEGFR2 (L-003148-00-0005) was incubated with Dharmafect 1 transfection reagent in Opti-MEM (51985034; Thermo Fisher Scientific) at RT for 30 min to allow complex formation. siControl, siEMCN, and hRECs were added in EGM-2 complete medium and plated on glass coverslips coated with 0.2% gelatin-PBS. Cells were incubated with transfection reagents for 48 hr before using in experiments.

### 2.8 Adenovirus infection

hRECs were seeded at 50% confluence 1 day prior to adenoviral infection. Cells were transduced with adenovirus expressing myc tagged human EMCN at a multiplicity of infection of 30 and cultured for 48 h inr EGM-2 medium supplemented with 2% FBS.

### 2.9 Immunoblotting

Cell lysates were collected using buffer containing protease inhibitors (Roche, Basel, Switzerland) and a phosphatase inhibitor cocktail (1:100, Sigma). Protein concentration was determined by the Bicinchoninic Acid Protein Assay (Thermo Scientific, Waltham, MA, USA, #23227). Proteins separated on SDS-PAGE gels were transferred to nitrocellulose membranes (VWR, #27376-991) and then probed with appropriate antibodies. Membranes were incubated with appropriate secondary antibodies and developed by fluorescence LI-COR Odyssey (LI-COR).

### 2.10 Immunofluorescence

Cells were fixed in 4% PFA for 10 min at room temperature and blocked in 3% BSA-PBS for 1 h at RT. The cells were then incubated in myc antibody overnight at 4 °C and secondary antibody for 1 h at RT. Following PBS washes, cells were permeabilized in 0.1% Triton-X 100 and incubated with the phalloidin-594 antibody for 30 min at RT. Cells were mounted using antifade reagent containing DAPI stain and images were taken using Zeiss Axioscope (Oberkochen, Germany).

### 2.11 Measurement of F/G-actin ratio

Intracellular G-actin and F-actin was separated using the G-Actin/F-Actin *In Vivo* Assay Biochem Kit (Cytoskeleton, BK037) according to the manufacturer’s instructions. Briefly, the cells were collected in lysis and F-actin stabilization buffer. Cell lysate was centrifugated at 350 x g for 5 min to remove cell debris. The supernatant was centrifuged at 100,000 x g for 1h (37°C) to separate the G-actin (soluble) and F-actin (insoluble). The supernatant and pellet which contain the G-actin and F-actin, respectively, were analyzed by immunoblotting.

### 2.12 Transmission electron microscopy

hRECs were grown on finder grids. Simultaneous extraction and fixation (0.25% Triton X-100; 0.25% glutaraldehyde) were used to preserve the actin cytoskeleton. Grids were then transferred to a 2% glutaraldehyde solution supplemented with 10 µg/mL phalloidin. Uranyl acetate was used for negative staining and images were collected using a FEI Tecnai G2 Spirit transmission electron microscope.

## 3. Results

### 3.1 Loss of EMCN reduces choroidal vascular sprouting

We examined whether the loss of EMCN affected choroidal angiogenesis using an ex vivo choroidal sprouting assay, which is used to study vessel growth in vitro.

Choroidal explants that were isolated from EMCN^flox/flox^ mice and cultured with complete medium for six days. The explants were incubated in culture medium containing adeno-associated virus (AAV)-Cre, which harbors Cre recombinase, or the AAV-EGFP control. Immunofluorescence staining revealed successful deletion EMCN (Fig. 1A). Sprouting was observed from days three to six. The sprouting area of EMCN deficient choroids (1.1 ± 0.2 mm^2^) was significantly reduced compared to that exposed to AAV-GFP control (2.6±0.4 mm^2^) (Fig. 1B-C).

**Figure 1.**
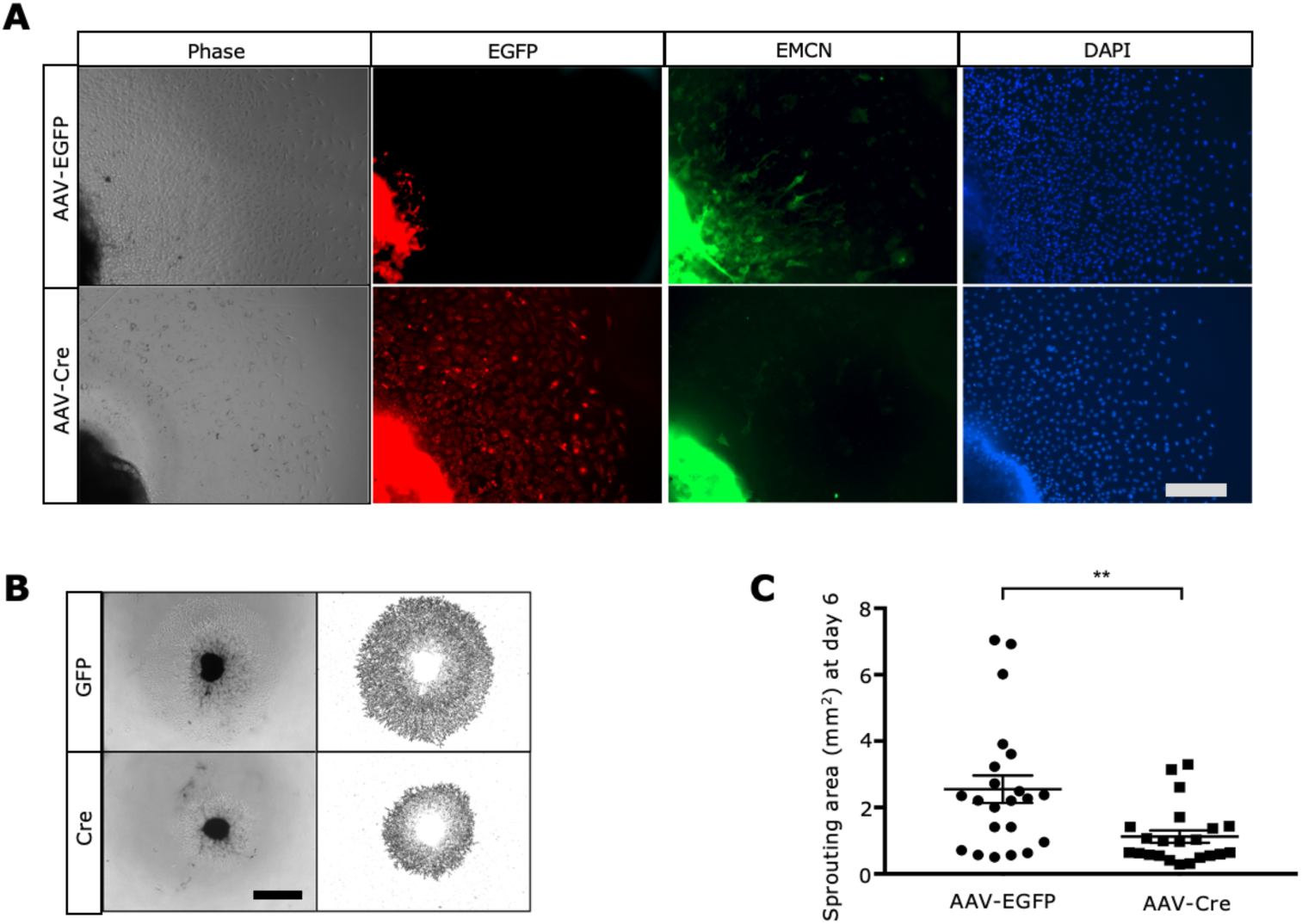
EMCN deletion significantly decreases choroid sprouting. Choroidal punches isolated from EMCN^flox/flox^ mice were cultured in Matrigel for 48 h and further cultured with AAV-EGFP or AAV-Cre recombinase to mediate EMCN deletion. (*A*) On day six, the samples were fixed and stained for EMCN (green). The GFP reporter gene (red) was switch on by Cre-mediated recombination. bar=500 µm. (*B-C*) The sprouting area was significantly reduced in choroidal explants with EMCN knockdown exposed to AAV-Cre (1.1 ± 0.2 mm^2^) compared to tissue treated with AAV-EGFP control (2.6±0.4 mm^2^). bar=1mm. \*\**p* < 0.005

### 3.2 Mass spectrometry identifies actin regulators interact with EMCN

Coordinated endothelial migration driving angiogenesis relies on changes in the actin cytoskeleton. Actin-binding proteins regulate the dynamics of the actin cytoskeleton in endothelial cells, thereby controlling cell-cell junction remodeling and migration. While the cytoskeletome of ECs has been widely investigated, the association between the glycocalyx and the actin cytoskeleton is not well characterized. We have previously observed that knockdown of EMCN resulted in fewer actin filaments in hRECs stimulated with VEGF compared to controls. To identify actin regulators that complex with EMCN in hRECs, we explored binding proteins by combining co-immunoprecipitation with mass spectrometry. The annotated proteins were analyzed using the PANTHER database. Fig. 2A-B display the Gene Ontology enrichment output and the protein classes, indicating that EMCN complexes with actin regulatory proteins. CFL2, ARPC3, ARPC1A, and TPM3 (Fig. 2C) were among the highly enriched proteins, and several of these proteins are known to play a conserved role in cell motility, contraction, and division.

**Figure 2.**
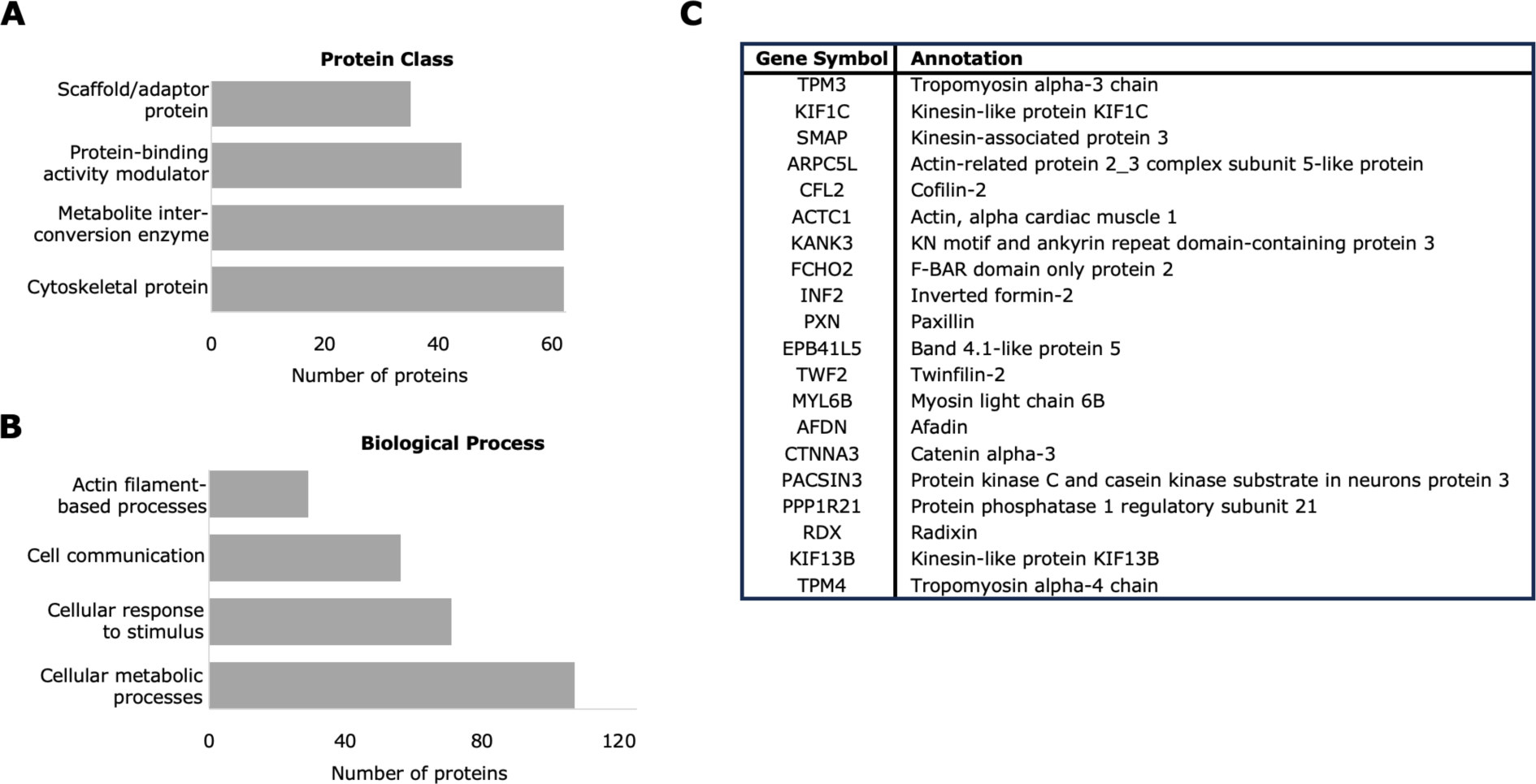
Proteins interacting with EMCN were identified via co-IP coupled with LC-MS/MS. 416 proteins that were annotated and further classified with respect to protein class (*A*) and biological process (*B*) using Protein ANalysis THrough Evolutionary Relationships (PANTHER). There were 56 cytoskeletal proteins, and 32 proteins that were annotated with actin filament-based processes. (*C*) The top 20 proteins in the cytoskeletal protein class.

### 3.3 EMCN expression modulates F-actin levels in hRECs

To gain insights into the involvement of EMCN in the regulation of the actin cytoskeleton, we knocked down or overexpressed EMCN by using siRNA or adenovirus, respectively, and stained with phalloidin. hRECs were harvested and analyzed 48 h following transfection with siControl or siEMCN (Fig. 3A). The depletion of EMCN resulted in reduced F-actin levels (Fig. 3B). In contrast, cells that were infected with an adenovirus overexpressing full-length human EMCN (hEMCN) displayed higher levels of F-actin. We observed that EMCN overexpression induced changes in organization of the actin filaments and cellular morphology. Compared to cells transduced with GFP adenovirus (control), hRECs overexpressing EMCN displayed a large number of cellular extensions and ruffles (Fig. 4A). The actin filaments, stabilized by supplemented phalloidin, were negatively stained and examined by TEM and revealed that hRECs overexpressing hEMCN contained of a dense actin network (Fig. 4B).

**Figure 3.**
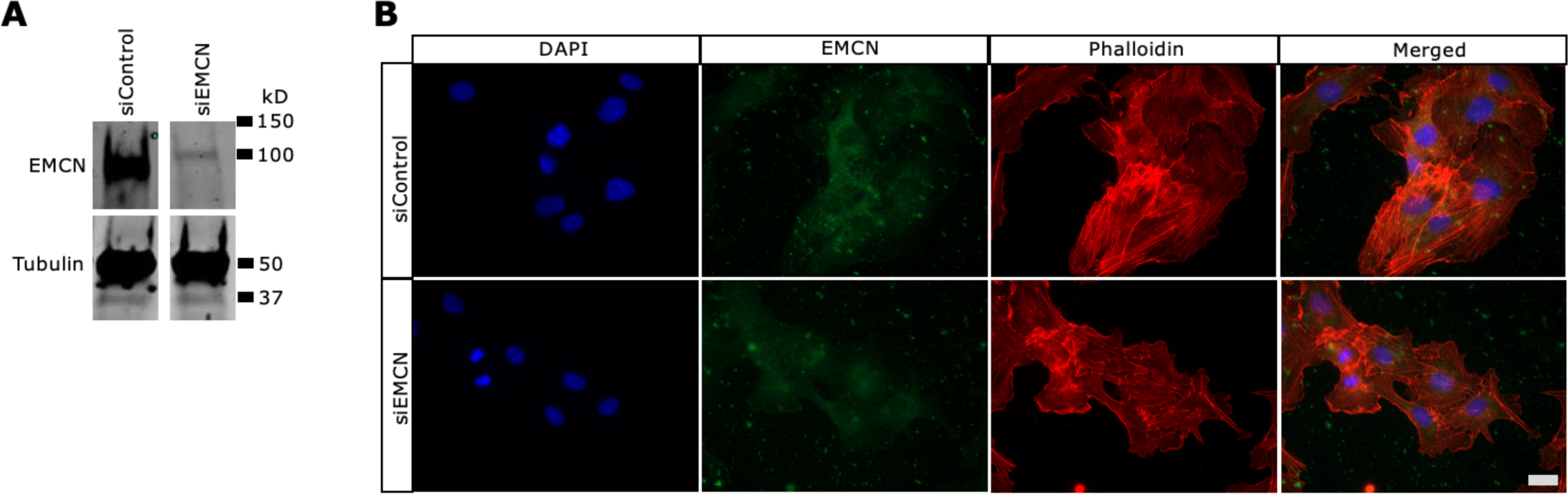
Reduced actin fibers with EMCN knockdown. HRECs were cultured in the presence of siControl or siEMCN (50 nM) for 48 h. (*A*) Knockdown of EMCN was verified by Western blot analysis. (*B*) Fixed cells were incubated with antibodies against EMCN (green) and phalloidin-594 (red). Silencing EMCN reduced stress fiber formation. bar=20 µm.

**Figure 4.**
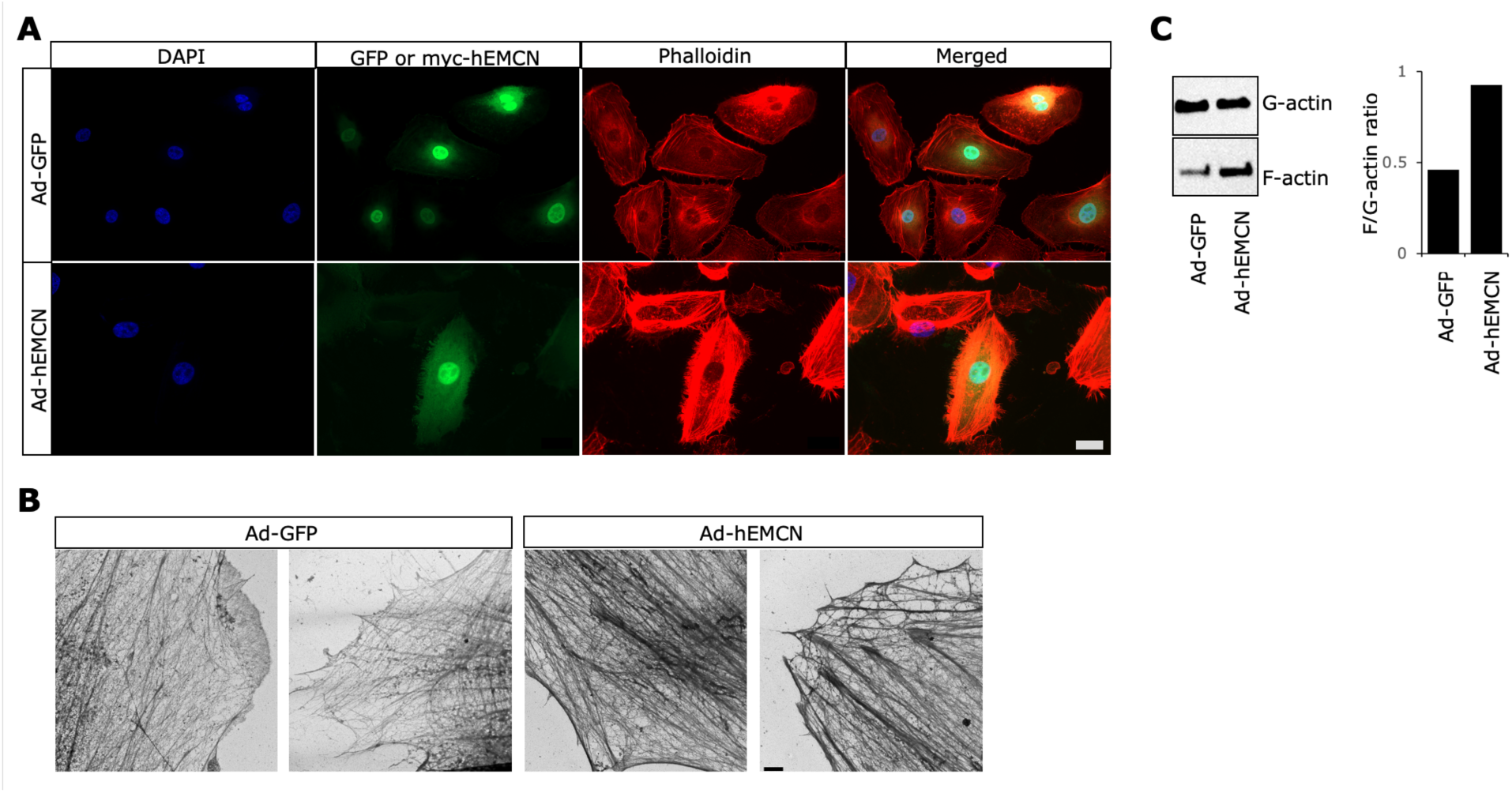
EMCN overexpression increases F-actin levels. Adenovirus expressing GFP (Ad-GFP) or human myc-tagged EMCN (Ad-hEMCN) (multiplicity of infection 30) were transduced into HRECs. (*A*) Cells were stained for EMCN (green) and F-actin (red). The HRECs overexpressing EMCN (green) had higher levels of F-actin (red), and multiple membrane protrusions compared to cells expressing GFP (control). bar=20 µm. (*B*) HRECs, cultured on gold electron microscopy grids and infected with Ad-hEMCN or Ad-GFP, were examined by FEI Tecnai G2 Spirit transmission electron microscope. Representative images of the cell edges show the dense network of filaments induced by EMCN overexpression. scale bar=2 µm. (*C*) Representative western blot of soluble G-actin and F-actin that was separated by centrifugation. There was a higher F/G-actin ratio in EMCN overexpressing cells.

Actin remodeling is a dynamic process of polymerization and depolymerization between the monomeric form (G-actin) and the filamentous polymer (F-actin). To assess the relative G-actin and F-actin content in hRECs overexpressing EMCN, the two actin pools were separated by centrifugation. The ratio of F/G-actin was higher in cells overexpressing EMCN (Fig. 4C).

### 3.4 The influence of EMCN on actin cytoskeletal dynamics is independent of VEGFR2

In EC, VEGF induces actin polymerization and the accumulation of stress fibers. Prior studies from our lab have highlighted that VEGFR2 internalization and downstream signaling, following VEGF stimulation, depend on its interaction with EMCN. To determine if the altered phenotype of EMCN overexpressing cells was an outcome of VEGF signaling, hRECs overexpressing EMCN were incubated with the VEGF trap aflibercept). Serving as a decoy receptor, aflibercept sequesters VEGF and prevents binding to the native receptor. There were no significant differences in the arrangement of the actin between cells treated with the VEGF inhibitor and those treated with PBS vehicle (Fig. 5). In another approach, endogenous VEGFR2 was knocked down using siRNA (Fig. 6A-B). Similar to the observation of VEGF neutralization, cells treated with siVEGFR2 group did not have any appreciable differences in F-actin levels and overall cell shape **c**ompared to cells treated with siControl (Fig. 6C).

**Figure 5.**
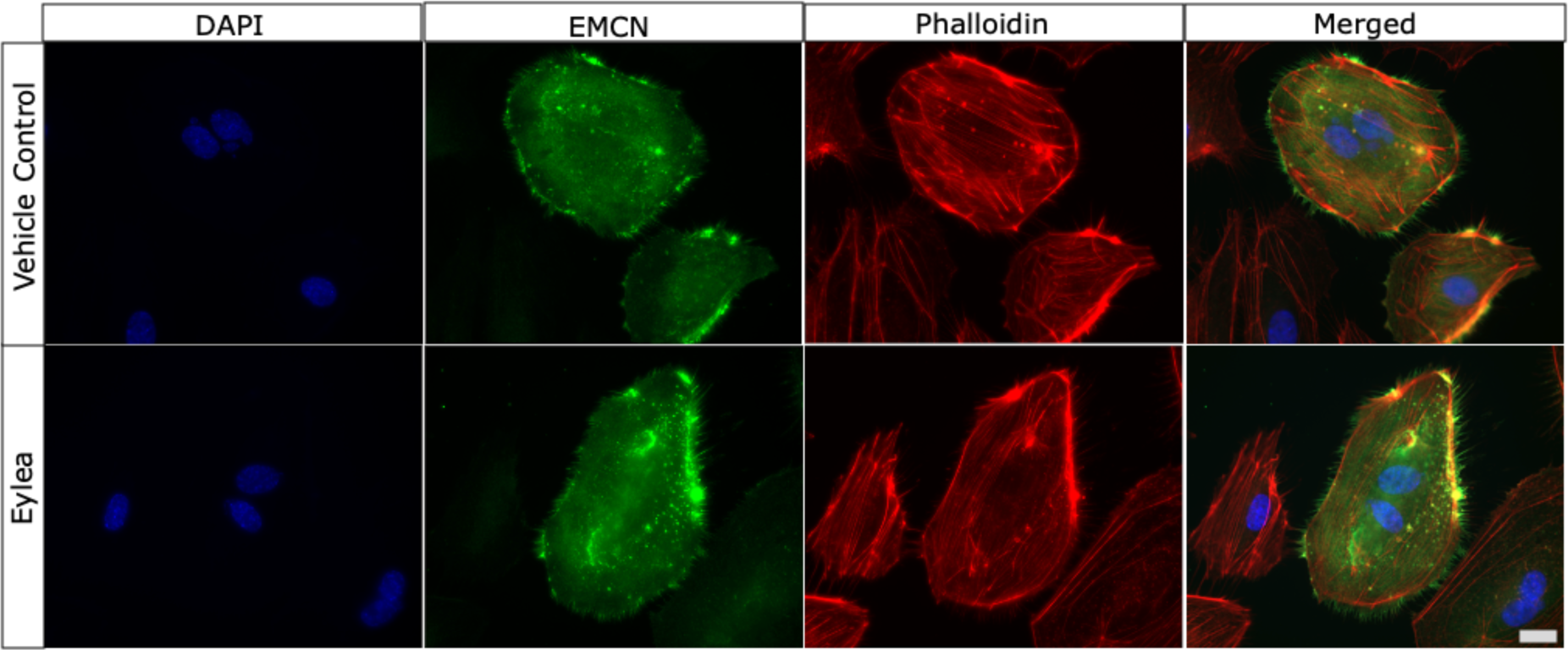
VEGF inhibition did not affect F-actin organization in EMCN overexpressing cells. HRECs that overexpress myc-tagged hEMCN were treated with aflibercept (1 nM) or PBS (vehicle) for 48 h. Representative immunocytochemistry images illustrate that changes in cell shape and actin reorganization persisted in the presence of VEGF inhibition. bar=20µm.

**Figure 6.**
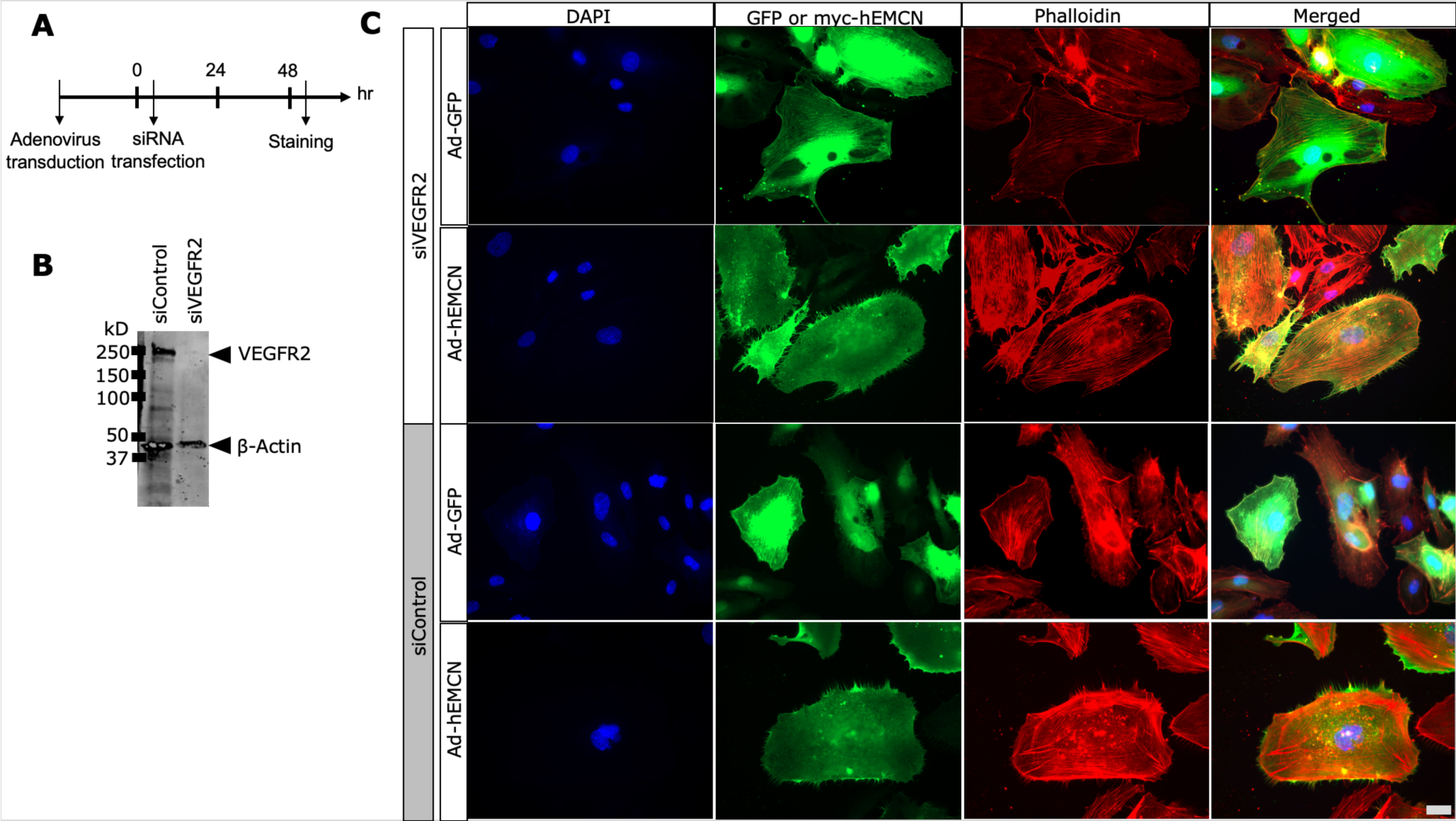
VEGFR2 signaling is not required for inducing cell morphology changes associated with EMCN overexpression. (*A*) 24 h after HRECs were transduced with Ad-hEMCN, endogenous VEGFR2 was knockdown using siRNA (50 nM). Cells were fixed and protein lysates were collected 48 h after siVEGFR2 treatment. (*B*) Representative immunoblot assessing efficiency of VEGFR2 knockdown. (*C*) No differences were observed in the GFP controls. Cells transduced with Ad-hEMCN underwent changes in the shape of the membrane. The absence of VEGFR2 did not affect the intense F-actin staining or cell morphology. bar=20µm.

## 4. Discussion

The actin cytoskeleton governs multiple cellular functions, and rapid remodeling of the endothelium is required for cell adhesion and migration [28, 29]. The glycocalyx is believed to be anchored to the actin cytoskeleton via core proteins such as syndecans, an interaction has been implicated in mechanosensation and transduction [30–32]. The mechanism by which the glycocalyx communicates changes to the cytoskeleton is less understood. VEGF/VEGFR2 signaling in endothelial cells activates downstream mechanotransductive signaling pathways and is the best-characterized axis in angiogenesis [33, 34]. EMCN is required for the VEGF stimulated internalization of the receptor and subsequent downstream signaling [16, 17]. Knockdown of EMCN not only reduces VEGF-induced cell proliferation but also results in less formation of stress fibers. The reduction in sprouting area of EMCN-depleted choroid biopsies supports the involvement of EMCN in angiogenesis. These effects suggest differential regulation of actin, which underlies the forces need to generate vessel sprouting. Here, we investigated the role of EMCN in the remodeling of the actin cytoskeleton.

Previous reports have demonstrated that glycoproteins expressed in glycocalyx are able to modify the shape of the plasma membrane [22–24]. The overexpression of the mucins caused membrane bending and induced membrane projections, likely as a result from steric hindrance and electrostatic repulsion between the side chains. These projections, or tubulations, were enriched in F-actin [24]. In our study, overexpression of EMCN in hRECs similarly generated spiky protrusions and resulted in increased F-actin levels. Interestingly, the actin was not localized solely to the cellular cortex, but there were also more transcytoplasmic actin stress fibers. Further classification of the actin stress fiber subtypes such as dorsal or ventral stress fibers and transverse arcs needs to be conducted. Notably, we found that dynamin was upregulated in hRECs overexpressing EMCN (unpublished). Apart from its role in endocytosis, dynamin colocalizes with actin filaments and align them into bundles in regions that undergo remodeling such as cortical ruffles [24, 35].

The increased actin polymerization that occurs in response to VEGF involves recruitment of the Nck adaptor protein to VEGFR2 [36]. Nck recruitment not only triggers the assembly of focal adhesions that enable actin bundling but also activates key regulators of actin nucleation such as actin Related Protein 2/3 complex (ARP2/3) [37–39]. To dissect whether the actin reorganization observed in hRECs overexpressing EMCN depends on VEGF signaling, we employed pharmacological inhibition of VEGF and genetic silencing of VEGFR2. In both cases, the cells remained rich in F-actin, suggesting that the changes in actin dynamics is VEGFR2-independent.

The cytoplasmic domain of syndecans has been proposed to directly or indirectly bind to the cortical actin network. Multiple actin binding proteins such as myosin II are engaged within approximately 100 nm in the submembrane space that collectively participate in the actomyosin contractility underlying membrane tension [32]. EMCN may also be interacting with other transmembrane proteins that recruit F-actin. Although it is possible that EMCN passively creates membrane projections, the high F-actin levels suggest continuous cycles of actin polymerization-depolymerization which requires ATP hydrolysis.

The actin cytoskeleton promotes structural stability of the endothelial glycocalyx under environmental factors including fluid shear stress [27, 40]. The actin cytoskeleton does not adapt once the glycocalyx is compromised, which leads to failure of cell migration and elongation. Components of the glycocalyx, including EMCN, affects actin reorganization, but whether the actin cytoskeleton has a reciprocal effect on the structure and function of the glycocalyx needs further examination.

## Notes

### Competing Interest Statement

The authors have declared no competing interest.

